# Functionally Essential and Structurally Diverse: Insights into the zebrafish Left-Right Organizer’s Cilia via Optogenetic IFT88 Perturbation and Volume Electron Microscopy

**DOI:** 10.1101/2025.09.08.674930

**Authors:** Favour Ononiwu, Melissa Mikolaj, Christopher Dell, Abdalla Wael Shamil, Kedar Narayan, Heidi Hehnly

## Abstract

In the zebrafish left-right organizer (LRO), the Kupffer’s Vesicle (KV), cilia extend from all cells into the fluid-filled lumen, but their structural diversity and contribution to morphogenesis remain incompletely defined. We hypothesized that cilia are required for KV development and may exist in distinct structural subtypes. Using a newly engineered transgenic line (*sox17:Cry2-GFP*), we optogenetically disrupted the intraflagellar transport protein IFT88 in KV progenitors via blue light-induced clustering of CIB1-RFP-IFT88. This perturbation impaired ciliogenesis and disrupted lumen formation, supporting a critical role for cilia in KV morphogenesis. To assess ciliary architecture, we used volume electron microscopy (vEM) to generate a high-resolution 3D ultrastructural map of mature KVs. Only 70.1% of cilia retained both mother and daughter centrioles, suggesting centriole elimination may occur in this tissue. Among centrioles present, 33.9% had distal appendages, 91.8% had subdistal appendages, and only 5.08% exhibited rootlet fibers. Cilia were also associated with membrane-bound vesicles, including spatially biased ciliary-associated vesicles (CaVs) and dense vesicles (CaDVs). These findings demonstrate that KV cilia are structurally diverse and spatially patterned, revealing a previously unappreciated level of complexity in LRO organization and providing new insight into how ciliary specialization may contribute to left-right axis specification.

## INTRODUCTION

The establishment of left–right (LR) asymmetry is a fundamental aspect of vertebrate development, essential for the proper positioning and morphogenesis of internal organs. In vertebrates, this symmetry-breaking process is initiated in the Left Right Organizer (LRO), a transient, ciliated organ. In zebrafish this organ is called the Kupffer’s Vesicle (KV) and is functionally analogous to the mammalian node. Motile cilia within the KV generate a unidirectional fluid flow that provides spatial cues necessary for activating LR patterning genes (Dasgupta and Amack, 2016; Grimes and Burdine, 2017). While cilia are clearly required for LR axis determination (Nonaka et al., 1998), the extent to which they contribute to the formation and organization of the LRO itself remains unresolved.

Despite progress in understanding cilia-driven flow (Layton, 1976; Lee and Anderson, 2008; McGrath et al., 2003; Nonaka et al., 1998; Okabe et al., 2008; Okada et al., 1999), surprisingly little is known about the role of cilia during early stages of LRO morphogenesis—when cells transition from progenitors to a structured cyst (for fish, (Essner et al., 2005)) or cup (for mice, (Lee and Anderson, 2008)) of cells surrounding a central fluid filled space. Whether cilia actively contribute to this morphogenic process or simply emerge as passive components once LRO architecture is established, is not known. An early study using a KIF3B mutant mouse demonstrated that LRO cells were unable to make cilia and had severe left-right developmental defects (Nonaka et al., 1998), but the morphology of the LRO itself was not robustly examined. Moreover, it remains unclear whether all cilia within the LRO are structurally equivalent or whether distinct ciliary subtypes exist with specialized functions.

Addressing these questions is critical to understanding how complex tissues form. Structural and functional heterogeneity among cilia could represent a previously unrecognized mechanism for fine-tuning organogenesis. To investigate this, we used a targeted optogenetic approach in zebrafish embryos to disrupt the function of the intraflagellar transport protein IFT88 with spatiotemporal precision, revealing a direct requirement for cilia in KV lumen development. In parallel, we performed 3D ultrastructural analysis using vEM to assess the diversity of cilia-associated structures within mature KVs. Together, these studies uncover a role for cilia in LRO morphogenesis and reveal unexpected structural heterogeneity, raising new questions about the cellular logic underlying ciliated tissue formation.

## RESULTS

### KV-specific optogenetic IFT88 clustering disrupts cilia formation and subsequent lumen development

To test the functional requirement of cilia during KV development, we developed a targeted optogenetic system to acutely and spatially disrupt intraflagellar transport within KV cells with the goal of disrupting cilia formation specifically. This system utilizes the CIB1/CRY2 blue light-inducible dimerization module (Aljiboury et al., 2023; Nguyen et al., 2016; Rathbun et al., 2020) to cluster and sequester the IFT-B complex component IFT88 (Pazour et al., 2000), thereby potentially inhibiting cilia assembly. Zebrafish embryos carrying the *Sox17:GFP-CAAX* transgene, which labels KV cells, were injected at the one-cell stage with mRNAs encoding CRY2 and/or CIB1-RFP-IFT88 causing global expression of both CRY2 and/or CIB1-RFP-IFT88 throughout the embryo. At 8 hours post-fertilization (hpf), embryos were exposed to blue light until 12 hpf, the developmental window during which KV forms and ciliogenesis is initiated (Figure 1A). Embryos were then fixed and immunostained for acetylated tubulin to mark cilia (Figure 1B, C). In controls expressing CRY2 alone, KV cells formed normal cilia. In contrast, co-expression of CIB1-RFP-IFT88 and CRY2 led to visible clustering of IFT88 and pronounced defects in cilia formation. Quantification across three independent clutches revealed a significant reduction in both the percentage of ciliated KV cells (Figure 1B, D) and cilia length (Figure 1E) in the IFT88-clustering condition compared to CRY2-only controls. Superplots were used for all analyses, displaying both individual embryo percentages or cilia lengths per embryo from each clutch and the corresponding means per clutch to visualize within- and between-experiment variability (Lord et al., 2020). These findings are consistent with previous data showing that CRY2 expression alone does not perturb ciliogenesis (Aljiboury et al., 2023), but when clustered with CIB1-RFP-IFT88 cilia formation is abnormal (Figure 1D, 1E). Together, these results demonstrate that IFT88 is required for KV ciliogenesis and establish optogenetic clustering as a precise and effective method for temporally controlled disruption of cilia formation in vivo.

**Figure 1.**
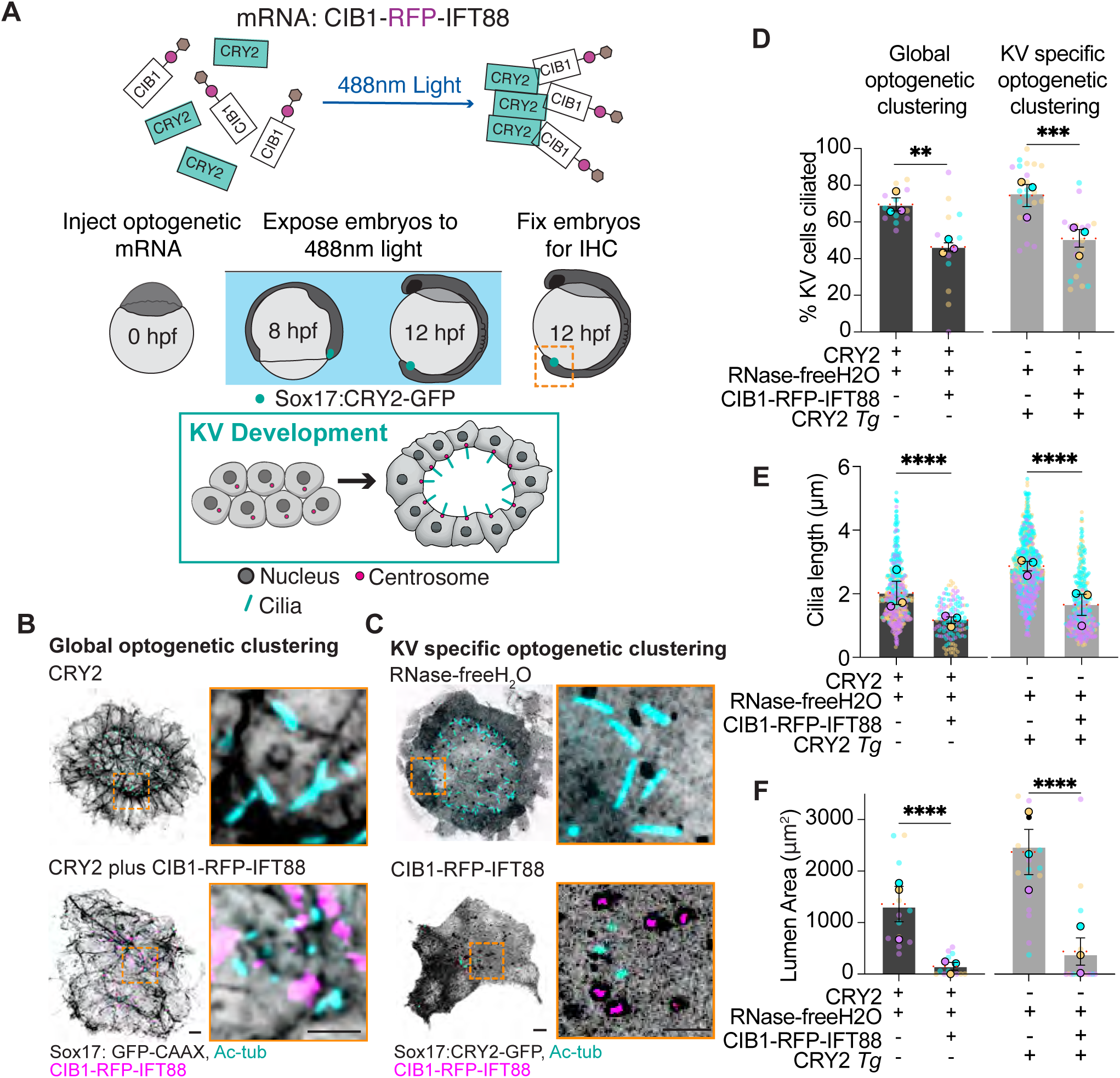
KV-specific optogenetic IFT88 clustering disrupts cilia formation and subsequent lumen development. **(A)** Schematic of the KV-specific optogenetic system. Embryos were injected at the 1-cell stage with mRNA encoding CIB1-RFP-IFT88, exposed to 488 nm light at 8 hpf to induce protein clustering, and fixed at 12 hpf. **(B, C)** Representative confocal images of *Sox17:GFP-CAAX* embryos injected with CRY2 and CIB1-mRuby-IFT88 mRNA, showing global (**B**) or KV-specific (**C**) optogenetic clustering. In (**B**), *Sox17:GFP-CAAX* marks KV plasma membranes (inverted grayscale); in (**C**), *Sox17:CRY2-GFP* labels KV cells (inverted grayscale). Acetylated tubulin (Ac-tub, cyan) labels cilia, and clustered IFT88 is visualized by CIB1-mRuby-IFT88 (magenta). Insets highlight clustered IFT88 (magenta) and KV cell membranes (inverted grayscale in **B**) or CRY2 expression (inverted grayscale in **C**). Scale bars, 10 µm (main), 5 µm (insets). **(D–F)** Bar graphs are superplots showing the percentage of KV ciliated cells (**D**), cilia length (**E**), and lumen area (**F**) under global and KV-specific clustering conditions. Data are mean ± SEM of clutch averages; each point represents an individual embryo color-coded by clutch (n=3). Statistical significance was determined by unpaired two-tailed t-tests (***p < 0.0001). Sample sizes per group: CRY2 control (n=13 embryos), global clustering (n=12), CRY2 transgenic control (n=22), KV-specific clustering (n=24). Refer to statistics Table S1.

To determine whether IFT88 function within KV cells is required for KV morphogenesis, we generated a *Sox17:CRY2-GFP* transgenic line, enabling spatially restricted optogenetic clustering of IFT88 specifically in KV cells. Embryos from this line were injected with *CIB1-RFP-IFT88* mRNA, and blue light exposure was applied from 8 to 12 hpf, corresponding to the developmental window of KV formation and ciliogenesis (Figure 1A). Consistent with our global clustering experiments using CRY2 and *CIB1-RFP-IFT88* mRNA, KV-specific clustering of IFT88 resulted in pronounced defects in ciliogenesis. Immunostaining for acetylated tubulin revealed a significant reduction in both the percentage of ciliated KV cells (Figure 1C, D) and cilia length (Figure 1E) compared to control *Sox17:CRY2-GFP* injected with RNase-free H_2_O. CIB1-RFP-IFT88 and CRY2 clusters were also observed at sites of shortened or stalled cilia that failed to elongate fully (Figure 1C). In addition to these ciliogenesis defects, KV-specific clustering of IFT88, similar to global disruption, led to a marked reduction in KV lumen area (Figure 1F), suggesting that cilia contribute not only to fluid flow but also to the morphogenetic processes that drive lumen formation. These results demonstrate that IFT88 activity within KV cells is essential for ciliogenesis and KV morphogenesis, supporting a model in which cilia play an active, cell-autonomous role in shaping the LRO.

### All KV cilia have γ-tubulin–positive centrosomes and IFT88, but Rootletin association is variable

To assess potential heterogeneity in ciliary morphology and molecular composition among KV cells, we first examined whether all KV cilia uniformly contain IFT88, a core component of intraflagellar transport, as a marker of conserved structure. In parallel, we evaluated the presence of Rootletin, a rootlet-associated component, to test whether rootlet structures are more variable across cilia. Ciliary rootlets are cytoskeletal structures that extend from the basal body into the cytoplasm and are primarily composed of the coiled-coil protein Rootletin (Mahen, 2021; van Hoorn and Carter, 2024). These striated filaments are well characterized in primary cilia, where they provide mechanical support and help anchor the basal body to the cytoskeleton (Antoniades et al., 2014; Lechtreck and Melkonian, 1998; Mahen, 2021; Soh et al., 2020). In cells with motile cilia, including multiciliated epithelia, rootlets have also been observed and are proposed to stabilize basal body positioning, coordinate basal body orientation, and resist mechanical strain during ciliary beating (Basquin et al., 2019; Frankel and Jenkins, 1979; Galati et al., 2014; Jerka-Dziadosz et al., 1995). Despite these proposed roles, the distribution and functional relevance of rootlets in the context of vertebrate left-right organizers, where motile cilia are essential for generating directional flow, remain unclear.

To investigate the spatial organization of the ciliary Rootlet component, Rootletin, in relation to the cilia and the centrosome in KV cells, we performed immunostaining at 12 hpf for acetylated tubulin (Ac-tub) to label cilia, γ-tubulin as a centrosome marker, IFT88 as a component of the intraflagellar transport complex that concentrates to the cilia, and Rootletin to mark ciliary rootlets (Figure 2A). Confocal imaging revealed that Ac-tub robustly labeled elongated cilia projecting from the apical surface of KV cells, while γ-tubulin localized to discrete puncta at the ciliary base (Figure 2B). IFT88 colocalized with Ac-tub along the axoneme and at the ciliary base, confirming its long-identified role in intraflagellar transport (Pazour et al., 2000) (Figure 2C). Rootletin was detected adjacent to γ-tubulin–positive centrosomes, consistent with its localization to the ciliary rootlet structure (Figure 2D), and was also observed as speckled puncta throughout KV cells.

**Figure 2.**
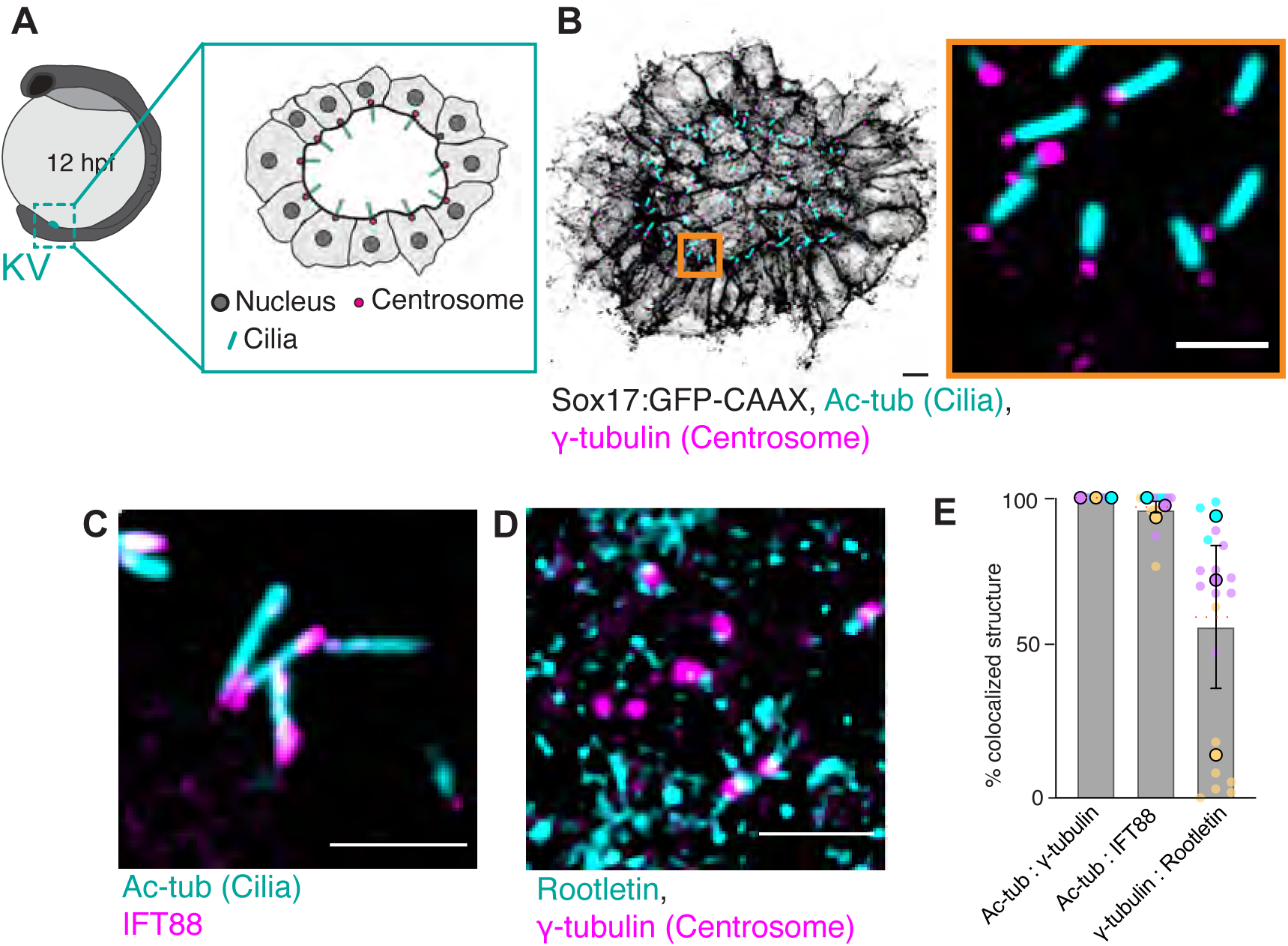
All KV cilia have γ-tubulin–positive centrosomes and IFT88, but Rootletin association is variable. **(A)** Schematic of a whole zebrafish embryo with an inset showing KV cells at 12 hpf. The inset illustrates the KV lumen surrounded by epithelial cells with nuclei (gray), centrosomes (red), and cilia (blue). **(B)** Confocal image of KV cells from *Sox17:GFP-CAAX* embryos (membranes, inverted grayscale) immunostained for acetylated tubulin (Ac-tub; cyan), marking cilia, and γ-tubulin (magenta), marking centrosomes. Inset shows a magnified view of cilia and centrosomes. Scale bars: 10 µm (main), 5 µm (inset). **(C-D**) Higher magnification images showing colocalization of acetylated tubulin (cyan) with IFT88 (magenta) along the cilium (**C**) or γ-tubulin (magenta) with Rootletin (cyan) at the centrosome (**D**). Scale bars, 5 µm. **(E)** Quantification of colocalization events: percentage of cilia (Ac-tub+) that also contain γ-tubulin (Ac-tub:γ-tubulin) or IFT88 (Ac-tub:IFT88), and percentage of γ-tubulin–positive centrosomes colocalized with Rootletin (γ-tubulin:Rootletin). Data represent mean ± SEM of individual embryos, with data points color-coded by clutch (n=3 clutches). Sample sizes: Ac-tub:γ-tubulin (n=29 embryos), Ac-tub:IFT88 (n=18), γ-tubulin:Rootletin (n=21). Refer to statistics Table S1.

To quantify the organization of these components, we assessed the association frequency across three independent clutches of embryos. We measured whether each ciliary structure (Ac-tub+) had associated γ-tubulin (Ac-tub:γ-tubulin) and IFT88 (Ac-tub:IFT88), and whether each γ-tubulin–positive centrosome was associated with Rootletin (γ-tubulin:Rootletin) (Figure 2E). As expected, all cilia were positive for both γ-tubulin and IFT88, with γ-tubulin localized at the ciliary base and IFT88 distributed at the base and along the axoneme. However, the proportion of γ-tubulin–positive centrosomes exhibiting detectable Rootletin signal varied across clutches, with a mean of 60.33 ± 23.78%, indicating heterogeneity in Rootletin centrosome recruitment and potential rootlet formation among KV cells. Notably, the presence of Rootletin alone is insufficient to confirm rootlet structures; confirmation would require electron microscopy. These findings suggest that KV cilia are compositionally heterogeneous, which may have functional consequences for ciliary stability or sensory capacity during KV morphogenesis.

### Workflow Overview for vEM Imaging of the Kupffer’s Vesicle KV

To investigate the ultrastructural heterogeneity of KV cilia, we employed vEM. vEM is a high-resolution imaging technique that combines serial ultrathin sectioning with Scanning Electron Microscopy (SEM) followed by computational reconstruction to generate three-dimensional nanoscale maps of cellular architecture. Unlike conventional confocal or super-resolution microscopy, which is limited by optical diffraction and antibody penetration, vEM enables the visualization of fine subcellular features such as basal bodies, rootlets, and ciliary membrane specializations with nanometer precision. This technique has been successfully applied to study complex tissue architectures and organelle organization in diverse systems, including cilia and centrosomes (Castranova et al., 2025; Cheng et al., 2019; Insinna et al., 2019; Micheva and Smith, 2007). By applying this approach to the zebrafish KV, we aimed to resolve previously uncharacterized features of centrosome and cilia architecture.

To characterize KV cilia at high resolution, we employed a serial-section vEM workflow. *Sox17:GFP-CAAX* transgenic zebrafish embryos at 12 hpf were first screened by fluorescence stereomicroscopy to identify fully formed GFP-positive KVs (Figure 3A). Embryos were then fixed, and the KV was micro dissected for downstream processing. Dissected KVs were mounted onto gridded coverslips (Figure 3B) to enable spatial tracking throughout the imaging pipeline.

**Figure 3.**
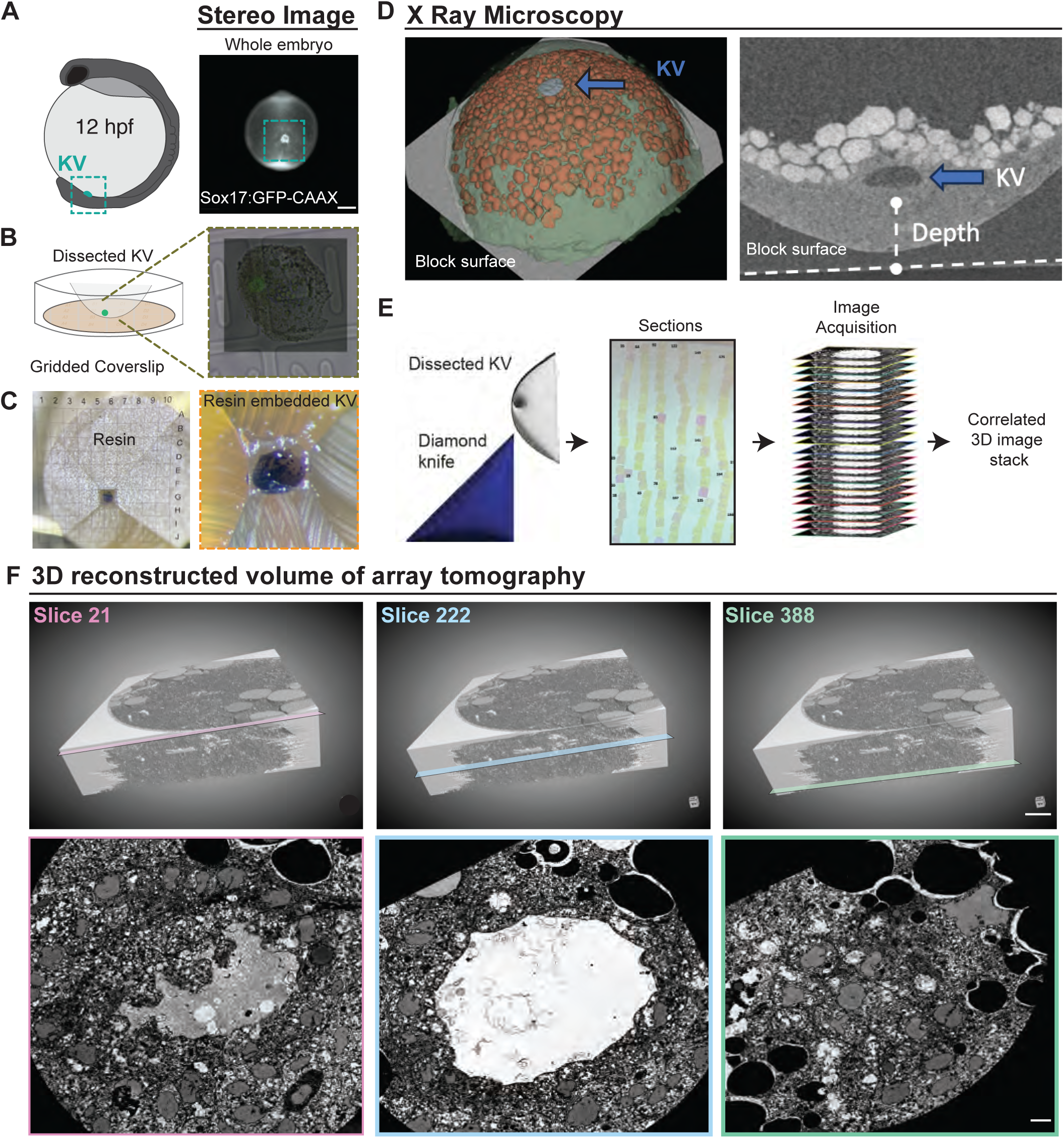
Workflow overview for vEM imaging of the Kupffer’s Vesicle (KV). (**A**) Model of a zebrafish 12 hpf embryo highlighting the KV (cyan, left), alongside an image of a real 12 hpf embryo with Sox17:GFP-CAAX–labeled KV (cyan box) acquired by stereomicroscopy (grey). Scale bar, 2 µm. (**B**) Schematic of dissected KV mounted on a gridded coverslip, inset shows corresponding microscope image. (**C**) Resin-embedded KV with gridded scale; zoomed image shows KV within the resin block. (**D**) X-ray microscopy image of the resin block with segmented annotations for skin (green), yolk cells (red), and KV (blue); X-ray image of the KV (inverted gray). (**E**) Schematic of ultramicrotomy process; schematic of computational stacking of acquired images. (**F**) Volumetric reconstruction of the KV with boxes indicating three slice locations; corresponding two-dimensional (2D) slices from specified serial sections. Scale bar, 5 µm.

The samples were embedded in resin with the KV positioned near the block surface to facilitate sectioning. Block face imaging relative to the grid confirmed correct positioning (Figure 3C). To ensure proper internal orientation, we performed X Ray microscopy. This technique is increasingly used to study mouse embryonic development, as it enables acquisition of high-contrast, high-resolution datasets of whole embryos (Handschuh and Glösmann, 2022). In our study, X Ray microscopy was used to segment major structures, including the embryonic cells (green), yolk cells (red), and KV (blue) (Figure 3D). These images confirmed the internal morphology of the KV and facilitated more accurate embryo alignment for optimal sectioning (Figure 3D). Ultrathin sectioning was performed using a diamond knife to generate ribbons of ∼70 nm sections (Figure 3E). These serial sections were imaged and computationally aligned to produce a volumetric reconstruction of the entire KV. Representative cross-sectional views from slices 21, 222, and 388 illustrate the well-preserved morphology and continuity of the KV structure (Figure 3F, Video S1). Owing to the time-intensive and technically demanding nature of this pipeline, from embryo preparation and X-ray microscopy to serial sectioning, imaging, and computational alignment, we successfully generated one complete volumetric dataset of an entire KV. All subsequent analyses are based on this single, high-quality dataset, highlighting both the strength and the current limitation of this approach.

### vEM of the zebrafish KV reveals that most cilia associate with both mother and daughter centrioles, but a subset lack one or both

To characterize the three-dimensional architecture of the mature KV, we performed high-resolution volumetric segmentation of the vEM dataset described in Figure 3. Image stacks were imported into Dragonfly (Object Research Systems), an advanced software platform for visualization, segmentation, and quantitative analysis of large-scale microscopy datasets (as in (D’Imprima et al., 2023)). Using Dragonfly, we reconstructed the entire KV volume and segmented key subcellular and tissue-level structures including cell nuclei, cilia, mother and daughter centrioles, the overall KV epithelium, and the central lumen (Figure 4A, Video S2). The resulting volumetric reconstruction revealed the mature KV dimensions—83 μm (x-axis), 79.5 μm (y-axis), and 40.7 μm (z-axis)—and permitted detailed spatial analysis of cilia positioning (Figure 4A). Nuclei were localized basally, while cilia projected apically into the lumen, consistent with the polarized architecture of KV cells (Figure 4B).

**Figure 4.**
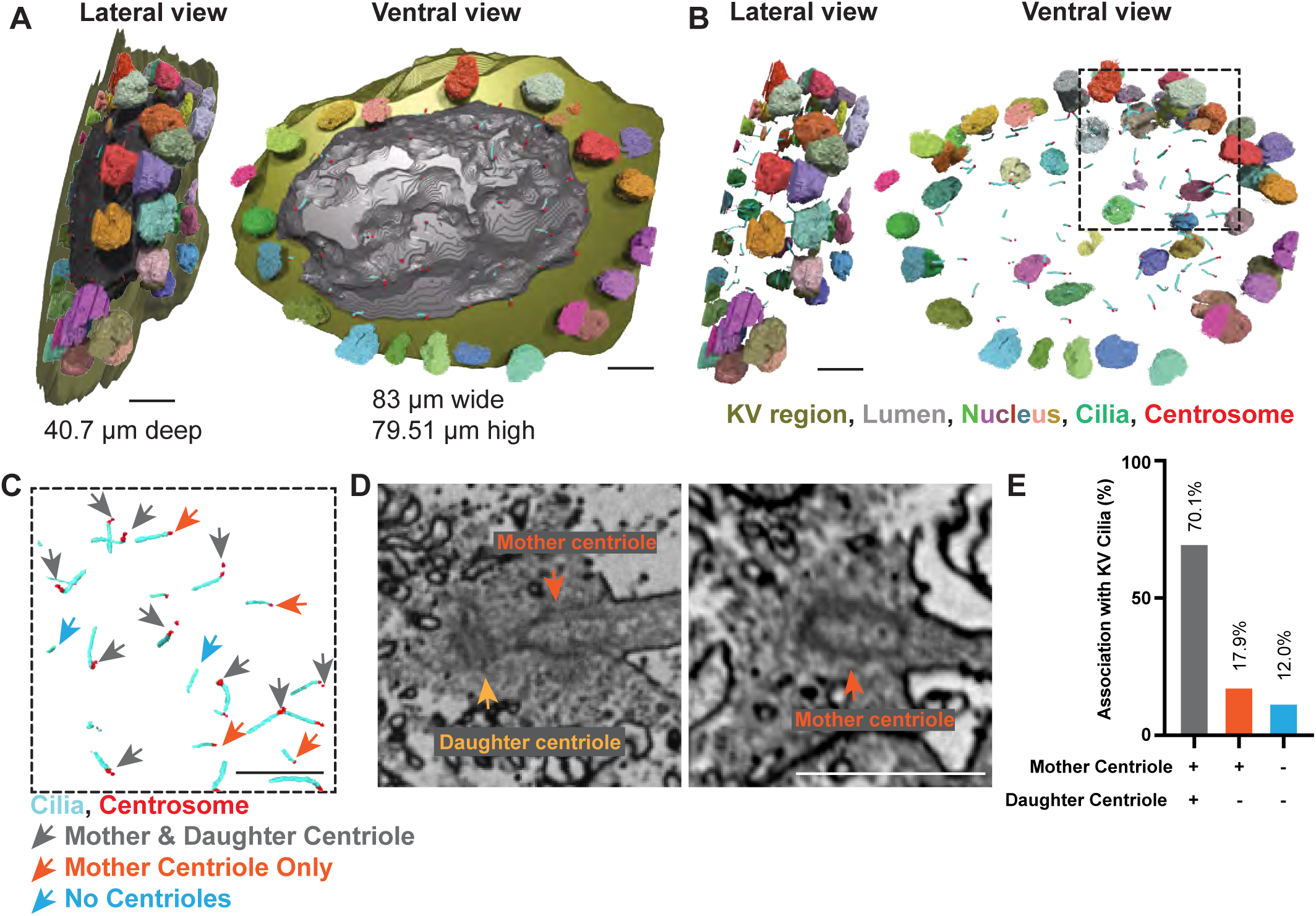
vEM of the zebrafish KV reveals that most cilia associate with both mother and daughter centrioles, but a subset lack one or both. **(A)** 3D reconstruction of the segmented KV volume showing overall dimensions. Left: Lateral view with KV region (green), lumen (gray), nuclei (multi-colored), cilia (cyan), centrosomes (orange), and z-axis measurement (40.7 µm). Right: Ventral view with x-axis (83 µm) and y-axis (79.51 µm) measurements. Scale bar, 10 μm. **(B, C)** 3D segmentation of KV organelle distribution showing nuclei (multi-colored), cilia (cyan), and centrosomes (orange). Left: Lateral view; Right: Ventral view. (**C,** inset) Zoomed view of segmented cilia and centrosomes showing cilia having mother centriole and daughter centriole (grey arrow), mother centriole only (orange arrow) and no centriole (blue arrow). Scale bar, 5 µm. **(D)** Single 2D slice from the array tomography dataset showing a KV cell with clearly resolved mother and daughter centrioles (Left) or mother centriole only (Right). Scale bar, 1 µm. **(E)** Bar graph quantifying the percentage of KV cilia in a single embryo that are associated with no centrioles, only a mother centriole, or both mother and daughter centrioles. Data were obtained from a fully segmented vEM dataset. (n=67 cilia). Refer to statistics Table S1.

Centriole number is typically tightly regulated. Cycling cells possess a pair of centrioles, a mature mother centriole, distinguished by the presence of distal and subdistal appendages and capable of templating a cilium, and a daughter centriole, which matures in the subsequent cell cycle (Vertii et al., 2016). In contrast, centriole number can be reduced in terminally differentiated cells, sometimes decreasing from two to one or even none. Such centriole elimination has been documented in the female and male germline, and in somatic cells during terminal differentiation, particularly in *C. elegans* and *Drosophila* (reviewed in (Kalbfuss and Gönczy, 2023)). While it is generally assumed that primary ciliated cells retain centrioles, with the mother centriole serving as the basal body for the ciliary axoneme, recent work in *C. elegans* has challenged this view. In sensory neurons, centrioles degenerate after initiating axoneme assembly (Serwas et al., 2017), leaving only focused pericentriolar material at the ciliary base (Garbrecht et al., 2021; Magescas et al., 2021), suggesting that even ciliated cells may eliminate centrioles under certain conditions.

To investigate whether KV cells retain both centrioles, we segmented and classified centrosomes based on the presence of mother and daughter centrioles (Figure 4C, D). Quantitative analysis of 67 segmented cilia revealed that 47 (70.1%) were associated with both mother and daughter centrioles, 12 (17.9%) with only a mother centriole, and 8 (12.0%) lacked any detectable centriole (Figure 4E, Video S2). Although centrioles are generally considered stable organelles that persist through multiple cell cycles, their selective elimination appears possible. While immunofluorescence in Figure 2 revealed that all cilia had γ-tubulin at their base, this approach does not provide sufficient resolution to determine whether the underlying centriole structure remains intact. Our vEM data resolve this ambiguity, revealing that even when centrioles are lost, centrosome-associated components such γ-tubulin likely persist. These findings highlight unexpected heterogeneity in centriole–cilia configurations among KV cells and provide a detailed ultrastructural map of centriolar architecture in the mature zebrafish left-right organizer.

### Structural heterogeneity of mother centrioles associated with KV cilia includes variable appendages and rootlet formation

To extend our analysis of ciliary and centrosome heterogeneity in KV cells, we examined centrosome-associated structures in the vEM dataset, building on earlier observations from Figures 2 and 4. We focused on rootlets and known features of the mother centriole, specifically distal appendages (DAs) and sub-distal appendages (SDAs) (modeled in Figure 5A), as well as additional electron-dense and membrane-associated elements. In most vertebrate cell culture systems, when cells are not ciliated, distal appendages (DAs) and sub-distal appendages (SDAs) are typically organized with ninefold symmetry. DAs are critical for docking the mother centriole to the plasma membrane, thereby licensing ciliogenesis, while SDAs serve as anchoring sites for microtubules and, in some ciliated cells, are reduced to a single structure known as the basal foot. The basal foot plays an essential role in orienting the cilium relative to the cell cortex and in coordinating tissue-level processes that depend on directional ciliary beating (reviewed in (Hall and Hehnly, 2021)). These observations suggest that appendage organization, while often stereotyped, may contribute an additional layer of heterogeneity to centrosome and cilia structure in developing tissues.

**Figure 5.**
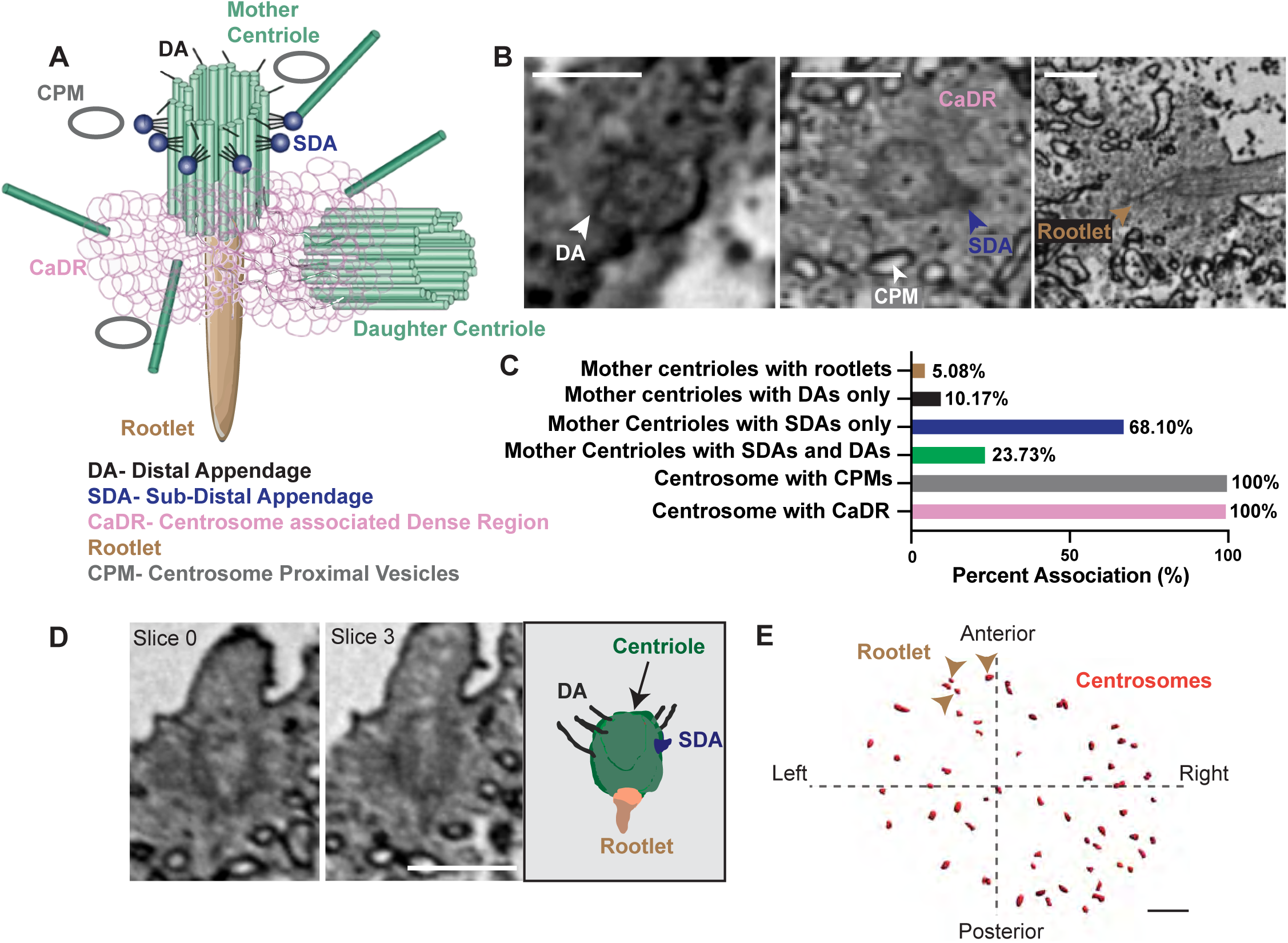
Structural heterogeneity of mother centrioles associated with KV cilia includes variable appendages and rootlet formation. **(A)** Schematic model of centrosome architecture highlighting the mother and daughter centrioles, distal appendages (DA, black), subdistal appendages (SDA, navy), centrosome-associated dense region (CaDR, pink), centrosome-proximal membrane (CPM, gray), and rootlet fibers (tan). Model adapted from (Doxsey, 2001). **(B)** Representative 2D single-slice images from the vEM dataset showing examples of centrosome-associated structures, including DAs, SDAs, CaDRs, CPMs, and rootlet fibers. Colored arrows and labels indicate each structure. Scale bar, 0.5 µm. **(C)** Bar graph quantifying the percentage of centrosomes with each associated structure among KV cells with mother centrioles (for rootlets, DAs, SDAs) or cilia (for CPMs, CaDRs). (n=59 mother centrioles). Refer to statistics Table S1. **(D)** Single 2D slices from the vEM dataset showing a KV mother centriole with resolved distal and subdistal appendages and a rootlet. A 3D segmented mother centriole (green) with distal appendages (DA, black), subdistal appendages (SDA, navy) and rootlet fibers (tan) from dataset on left is shown. Scale bar, 1 µm. **(E)** 3D reconstruction of the KV showing the spatial distribution of centrosomes (red dots) and centrosomes with rootlet fibers (tan arrows) along the anterior–posterior axis of the embryo. Scale bar, 5 µm.

During segmentation and analysis, we consistently observed a Centrosome-associated Dense Region (CaDR) surrounding centrioles, which may correspond to the pericentriolar material (PCM) ((Doxsey et al., 1994), Figure 5A), and centrosome-proximal membranes (CPMs) (Figure 5A), likely representing centriolar vesicles described in previous studies (H. Hehnly, C-T. Chen, C. Powers, H-L Liu, 2012; Knödler et al., 2010; Westlake et al., 2011). The mother and daughter centrioles were readily distinguishable (Figure 5B), with appendages localized exclusively to the mother centriole. CaDRs and CPMs were consistently present near centrioles across all cells examined (Figure 5B). Rootlet fibers were also visualized (Figure 5B).

To quantify the prevalence of centrosome-associated features, we analyzed their presence across the entire segmented KV volume (Figure 5C). Centrosome-associated dense regions (CaDRs) and centrosome-proximal membranes (CPMs) were consistently observed in 100% of cilia, including the 12% of cases lacking detectable centrioles, suggesting that these structures may persist beyond centriole loss and play a fundamental role in maintaining ciliary function. In contrast, centriole appendage composition was more variable: 23.7% of mother centrioles contained both distal appendages (DAs) and sub-distal appendages (SDAs), 68.1% had only SDAs, and 10.2% had only DAs. Overall, SDAs were observed more frequently (91.8%) than DAs (33.9%). While DAs consistently exhibited ninefold symmetry, SDAs rarely did (with only a single SDA often present; see Figure 5B, 5D). Notably, centrioles that were associated with rootlets (only 5.08%) always possessed both DAs and SDAs, suggesting that full appendage complement may be required for rootlet anchoring (3D segmentation of mother centriole containing a rootlet, DA, and SDA, Figure 5D). It is also possible that the visualization of DAs is underestimated due to their small size and orientation-dependent detectability in serial sections.

Despite this possible caveat, the data reveal notable heterogeneity in centriole architecture, with a subset of centrioles lacking SDAs entirely. This variability, along with the persistence of CaDRs and CPMs in the absence of centrioles, raises the possibility that KV cells may undergo progressive centriole elimination, as has been observed in other developmental contexts (Kalbfuss and Gönczy, 2023). Supporting this idea, rootlet fibers—typically associated with the mother centriole—were observed in a minority of KV cells and detected in only 5.08% of mother centrioles by structural analysis (Figure 5C). Interestingly, this contrasts with our immunostaining data, where a range of centrosomes (60.33 ± 23.78%) were positive for the rootlet-associated protein Rootletin (Figure 2). This discrepancy suggests that while Rootletin remains localized to the centrosome, it does not always assemble into a bona fide rootlet structure. Spatial analysis of the vEM data set revealed that rootlet-positive cilia were predominantly localized to the anterior-left quadrant of the KV (Figure 5E, 3 arrows). This asymmetrical enrichment suggests that ciliary composition varies regionally and may reflect distinct structural or mechanical roles for anterior-left cilia in generating or sensing fluid flow during left-right axis specification. Moreover, the observed heterogeneity in centrosome architecture may also reflect temporal dynamics, where some centrosomes have already disassembled their appendages or centrioles, whereas others have not yet initiated this process, thereby contributing to the spectrum of structures detected. However, unlike rootlets, centrioles and their associated appendages show no consistent spatial pattern of loss within the KV.

### Ciliary heterogeneity in KV cells is defined by a predominant cilia subtype exhibiting membrane-associated vesicles and lacking a ciliary pocket

While analyzing cilia in the vEM dataset, we identified cilia with discernible axonemes (modeled in Figure 6A and shown in Figure 6B). Motile cilia are classically described as having a 9+2 microtubule arrangement, whereas non-motile (sensory) cilia in vertebrates are primarily 9+0 (Marra et al., 2016). We had hoped to map the distribution of 9+0 and 9+2 cilia across KV cells; however, although some cross-sections (e.g., Figure 6B) suggest a 9+2 morphology, the resolution and orientation of many sections prevented confident assignment of axonemal architecture in all cases.

**Figure 6.**
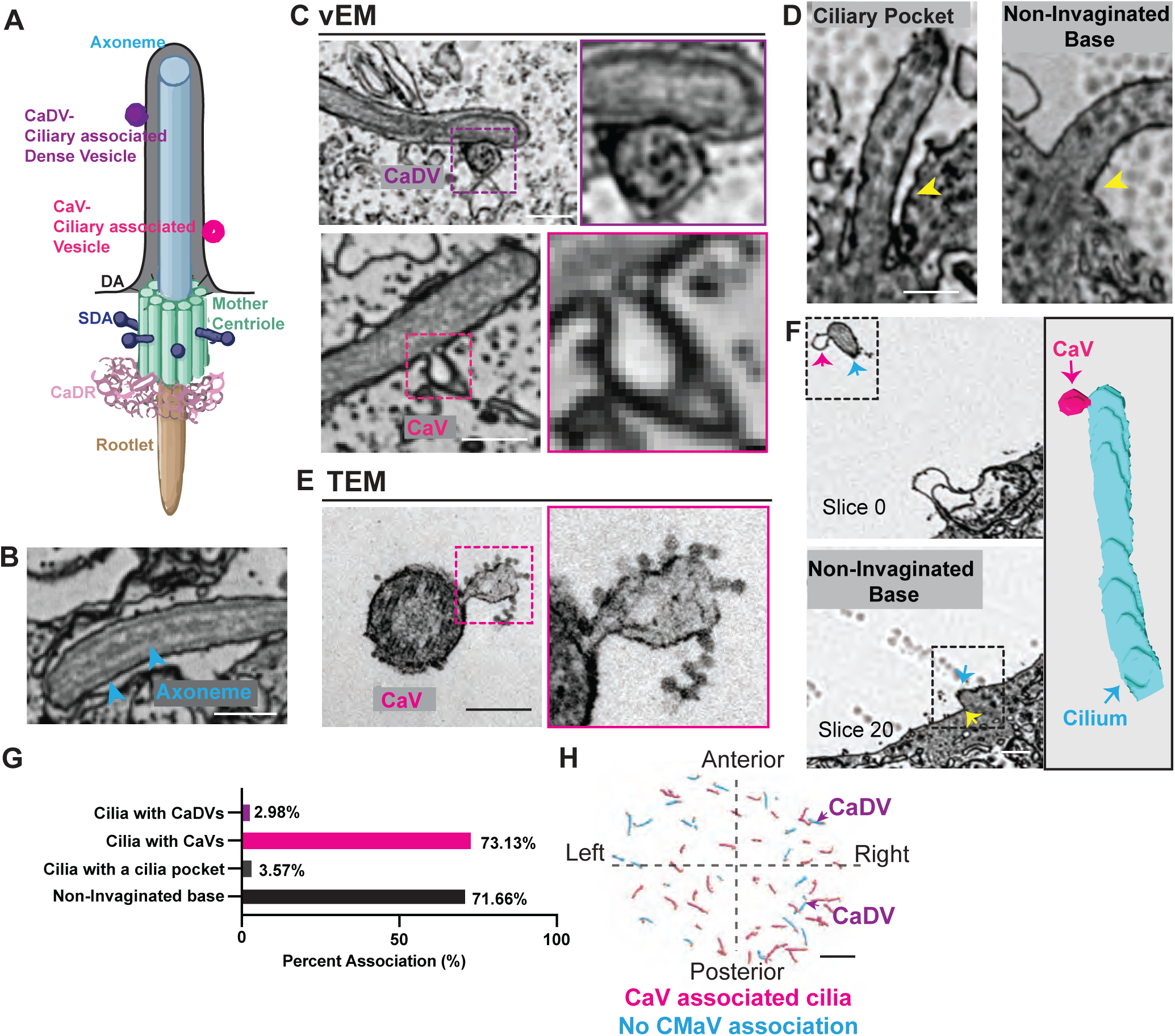
Ciliary heterogeneity in KV cells is defined by a predominant cilia subtype that exhibits membrane-associated vesicles and lacks a ciliary pocket. **(A)** Schematic representation of a cilium with an axoneme surrounded by a membrane bearing two types of associated vesicles: Ciliary-associated Dense Vesicle (CaDV, purple) and Ciliary-associated Vesicle (CaV, magenta). The mother centriole is depicted with distal appendages (DAs), subdistal appendages (SDAs), centrosome-associated dense region (CaDR), and rootlet. **(B–D)** Representative 2D slices from the serial vEM dataset. (**B**) Axoneme shown. (**C**) Cilia with CaDV (top) and CaV (bottom); insets show magnified views highlighting differences in vesicle morphology. (**D**) Examples of invaginated vesicle bases (IVB) and non-invaginated vesicle bases (NIVB). Scale bar, 0.5 µm. **(E)** Transmission Electron Micrograph shown of KV cilia cross-section that contains a ciliary vesicle. Scale bar, 0.2 µm. **(F)** Representative vEM slices from slice 0 to slice 20 (note the non-invaginated base), used for 3D reconstruction (right). The reconstruction demonstrates a KV cilium (blue) associated with a CaV (magenta arrow). Scale bar, 0.5 µm. **(G)** Bar graph quantifying the percentage of cilia associated with CaDV, CaV, IVB, and NIVB from the vEM dataset (n=67 cilia from KV dataset). Refer to statistics Table S1. **(H)** Spatial map of CaV+ (pink) and CaV– (blue) vesicles in KV cells, showing localization relative to the anterior–posterior and left–right axes of the embryo. Cilia associated with CaDV are indicated by purple arrows. Scale bar, 5 µm.

In contrast, what we could consistently resolve were cilia frequently accompanied by cilia-associated vesicles (CaVs), either fused to or positioned in close proximity to the axoneme (modeled in Figure 6A, shown in vEM images in Figure 6C, bottom, 3D segmentation of a cilium with CaV in Figure 6F). These vesicles often appeared to be forming from or fusing with the ciliary membrane, a process more clearly visualized by TEM (Figure 6E), suggesting that they may arise directly from the ciliary membrane or represent incoming vesicles from another source. By comparison, ciliary-derived dense vesicles (CaDVs) were observed only in association with the ciliary membrane, without discernible evidence of membrane fusion or pinching events at this resolution (Figure 6C, top), raising the possibility that these vesicles do not originate from the cilia but rather from an external source.

Ciliary membrane–associated vesicles have been reported in multiple systems (Nager et al., 2017; Nikonorova et al., 2025; Volz et al., 2021; Wang et al., 2021; Wang et al., 2024a; Wood et al., 2013), but to our knowledge have not been described in the LRO. These vesicles are thought to mediate the targeted delivery and removal of membrane proteins, reflecting active remodeling of the ciliary membrane. They have also been implicated in intercellular communication and the compartmentalization of signaling activity (Luxmi and King, 2022; Wang and Barr, 2016; Wang et al., 2024b).

Notably, we observed very few cilia displaying a ciliary pocket in this dataset (Figure 6D, F). The ciliary pocket is a membrane invagination at the base of the cilium implicated in regulating vesicular trafficking, selective cargo entry, and protein turnover during ciliogenesis. It is a hallmark feature of many vertebrate cell types and plays critical roles in sensory transduction and signal coordination (Ghossoub et al., 2011; Molla-Herman et al., 2010; Rohatgi and Snell, 2010).

Quantitative analysis revealed that most cilia exhibited membrane-associated structures, such as cilia-associated vesicles (CaVs; 73.13%). In contrast, only a small subset displayed cilia membrane-associated dense vesicles (CaDVs; 2.98%) or invaginated bases (IVBs/cilia pockets; 3.57%) (Figure 6G). Observations in which the presence or absence of a ciliary pocket could not be clearly resolved were excluded from the analysis. Spatial mapping showed that CaVs were more frequently observed on the posterior and left sides of the Kupffer’s Vesicle (KV), whereas CaDVs were restricted to the right side (cilia with CADVs, arrows, Figure 6H), supporting the existence of regionally specialized cilia-associated vesicle populations within the KV.

## DISCUSSION

This study reveals that cilia play a direct, cell-autonomous role in shaping KV morphogenesis. Using a KV-specific optogenetic system to acutely cluster IFT88, a core component of the IFT machinery (Pazour et al., 2000), we showed that disrupting cilia formation results in reduced lumen size and shortened cilia. This approach provides spatial and temporal precision beyond traditional knockouts, enabling a more refined dissection of cilia disruption within a discrete time frame and how this effects KV development (Figure 1). While our findings are consistent with IFT88’s established role in ciliogenesis (Pazour et al., 2000), they also raise the possibility that defects in lumen formation involve both cilia-dependent and potentially cilia-independent functions, as IFT proteins have been implicated in non-ciliary roles such as cell division and cell migration (Boehlke et al., 2015; Borovina and Ciruna, 2013; Delaval et al., 2011; Vertii et al., 2015; Wang et al., 2016). Distinguishing between these mechanisms will require further targeted manipulations and live imaging during KV assembly.

Beyond functional perturbation, our volumetric ultrastructural analyses revealed previously unrecognized diversity in ciliary architecture. Using vEM, we generated a comprehensive 3D reconstruction of the KV, segmenting cilia, nuclei, centrosomes, and the lumen. This revealed significant heterogeneity in centriole number and in the presence of associated structural features, including rootlets, DAs, and SDAs (Figure 5). Notably, while centrioles were consistently accompanied by centrosome-associated dense regions and proximal membranes, their appendage composition varied widely. SDAs were the most retained structure, followed by DAs and then rootlets, which were detected in only a minority of KV cells. This graded pattern raises the intriguing possibility that centriole elimination proceeds in a stepwise manner, beginning with loss of rootlet fibers, followed by disassembly of distal appendages, and eventually removal of subdistal appendages. Such a model is consistent with the observed abundance hierarchy of these structures. Analogous centriole remodeling events have been described in multiple systems. In *C. elegans* sensory neurons, centrioles degenerate following cilium formation, leaving only residual pericentriolar material at the base of the cilium (Garbrecht et al., 2021; Magescas et al., 2021; Serwas et al., 2017). Similarly, in vertebrate spermatids and oocytes, centriole loss occurs through progressive disassembly of structural components (Aljiboury and Hehnly, 2023). These findings suggest that centriole elimination can be a regulated, multistep process, and our data raise the possibility that similar mechanisms may operate within the KV to generate ciliary heterogeneity during left-right axis specification.

In addition to centriolar features, we identified two distinct classes of ciliary membrane-associated vesicles: ciliary-associated vesicles (CaVs) and ciliary-associated dense vesicles (CaDVs), with CaVs representing the predominant subtype (Figure 6). Notably, KV cilia lack a classical ciliary pocket. However, CaVs were frequently observed adjacent to the axoneme and non-invaginated base, suggesting that these vesicles may function as trafficking intermediates for membrane or protein delivery. Spatial mapping revealed an asymmetry in vesicle localization, with CaVs enriched on the posterior-left side of the KV and CaDVs restricted to the right side. This polarized distribution parallels the established axis of KV fluid flow and raises the possibility that region-specific vesicle populations contribute to spatial regulation of membrane dynamics or signaling events required for left-right axis specification.

In summary, our combined optogenetic and ultrastructural analysis provides new insight into the cellular mechanisms underlying KV morphogenesis and ciliary specialization. The approach establishes a framework for dissecting how cytoskeletal and membrane-based systems contribute to left–right organizer function and raises new questions about how structural diversity among cilia supports developmental symmetry breaking.

## Acknowledgements

We thank In Vivo Biosystems for helping us create the *Sox17:CRY2-GFP* line, Jesse Aaron and the Advanced Imaging Center (AIC), HHMI Janelia Research Campus for access to the Zeiss Versa XRM instrument for sample location within the resin block for subsequent vEM. Benjamin Zink at SUNY Upstate Medical University for performing the TEM imaging as a paid service.

## Author Contribution

F.O. designed, performed, and analyzed experiments, and wrote the first draft of the manuscript; H.H. oversaw the project, edited and wrote the manuscript; M.M., C.D., A.W.S. and K.N. performed experiments and associated analysis and edited the manuscript.

---

**Video S1.** *Workflow overview for vEM imaging of the Kupffer’s Vesicle (KV).* Video shows a series of 406 vEM slices through the zebrafish KV.

**Video S2.** *vEM of the zebrafish KV reveals that most cilia associate with both mother and daughter centrioles, but a subset lack one or both.* Video shows a 3D segmentation of entire KV. KV region (green), lumen (gray), nuclei (multi-colored), cilia (cyan), centrosomes (orange).

## Methods

### Zebrafish Lines

Zebrafish lines were maintained according to the approved protocols of the Institutional Animal Care Committee of Syracuse University (IACUC Protocol #18-006). Embryos were raised at 28.5°C and staged (as described in (Kimmel et al., 1995)). Transgenic zebrafish lines used for vEM and immunohistochemistry are listed in Key Resource Table S2.

### Plasmids and mRNA for injection experiments

Plasmids were generated using Gibson cloning methods (NEBuilder HiFi DNA assembly Cloning Kit) and maxi-prepped before injection and/or transfection. mRNA was made using mMESSAGE mMACHINETMSP6 transcription kit and mMESSAGE mMACHINETMT3 transcription kit. See the key resource table for a list of plasmid constructs and mRNA used. Injections of zygote stage embryos were performed as described in (Aljiboury et al., 2021).

### Optogenetic experiments in zebrafish embryos

At the zygote to one-cell stage, zebrafish embryos were injected with 50–200 pg of CRY2 and/or CIB1-RFP-IFT88. After injection, embryos were kept in darkness to develop until the 50% epiboly stage, when they were exposed to 488 nm light using the NIGHTSEA fluorescence system or the stereomicroscope. *Tg(Sox17: GFP-CAAX), TgBAC(cftr-GFP),* and *Tg(Sox17: GFP)* were used for global activation, while *Tg(Sox17:CRY2-GFP)* enabled KV-specific activation. Illumination continued until the six-somite stage (Rathbun et al., 2020). Following light exposure, embryos were fixed in 4% paraformaldehyde and immunostained according to established protocols (Aljiboury et al., 2021).

### Immunofluorescence

Immunostaining for acetylated tubulin, γ-tubulin, IFT88, Rootletin, and GFP was performed on transgenic zebrafish embryos [*Tg(Sox17:GFP-CAAX), TgBAC(cftr-GFP), Tg(Sox17:GFP), Tg(Sox17:CRY2-GFP)*] fixed at 8, 10, and 12 hours post-fertilization (hpf) in 4% paraformaldehyde (PFA) with 0.5% Triton X-100 at 4°C overnight. Embryos were dechorinated after washing with PBST (0.1% Tween-20 in Phosphate Buffered Saline (PBS)) 3 times. Embryos were blocked in wash solution (1%DMSO,1% BSA,0.1%Triton-X) for 4 hours at room temperature with gentle agitation. Primary antibody incubation (diluted in wash solution) occurs overnight at 4°C. Primary antibodies used include: anti-IFT88 (Rabbit) antibody (1:200, Proteintech, 13967-1-AP: AB_2121979), anti-γ-Tubulin (Goat) antibody (1:200, Santa Cruz, sc-7396), Anti-GFP (Chicken) (1:300, GeneTex, GTX13970: AB_371416), Anti-Acetylated Tubulin (Mouse) (1:300, Sigma Aldrich, T6793: RRID: AB_477585), Anti-GFP (Rabbit) (1:300, Molecular Probes, A-11122: AB_221569), and Anti-Rootlettin (Chicken) antibody (1:300, Fisher Scientific, ABN1686MI), refer to Key Resource Table S2. Embryos were washed, blocked for an hour, and incubated with secondary antibodies for 2-4 hours at room temperature or overnight at 4°C. Secondary antibodies used include: Alexa Fluor Anti-Mouse 568 (1:300, Life Technologies, A10037; RRID: AB_2534013), Alexa Fluor Anti-Chicken 488 (1:300, Fisher Scientific, A11039), or Alexa Fluor Anti-Mouse 647 (1:300, Life Technologies, A31571; RRID: AB_162542), Alexa Fluor Anti-Rabbit 568 (1:300, Life Technologies, A21206; RRID: AB_2535792), Alexa Fluor Anti-Rabbit 647 (1:300, Fisher Scientific, A31573; RRID: AB_2536183), Alexa Fluor Anti-Goat 647 (1:300, Jackson ImmunoResearch Laboraties, 705-605-003; RRID: AB_2340436). Embryos were stained with DAPI (1 mg/mL) to label nuclei after washing 3 times with wash solution. Embryos were mounted with 2% agarose after washing with PBS. Refer to Key Resource Table S2.

### Imaging

The Leica SP8 laser scanning confocal microscope (LSCM) and the Leica DMi8 with a spinning disk confocal were used to image KV development and lumen formation in zebrafish embryos. The SP8 system was equipped with HC PL APO 20x/0.75 IMM CORR, HC PL APO 40x/1.10 W CORR water, and HC PL APO 63x/1.3 Glyc CORR glycerol objectives, with image acquisition performed using LAS-X software. The DMi8 spinning disk system included an X-light V2 confocal unit, Visitron VisiFRAP-DC photokinetics (405 and 355 nm lasers), a Lumencore SPECTRA X light source, a Photometrics Prime-95B sCMOS camera, and an 89 North-LDi laser launch. Objectives used with this system included HC PL APO 40x/1.10 W CORR water, HC PL APO 40x/0.95 NA CORR dry, and HCX PL APO 63x/1.40–0.60 NA oil. Images were captured using VisiView software. For staging, a Leica M165 FC stereomicroscope equipped with a DFC 9000 GT sCMOS camera was used.

### Association analysis

Association between Ac-tubulin and γ-tubulin, Ac-tubulin and IFT88, and Rootletin and γ-tubulin signals was assessed by manually counting the number of cilia or centrosomes exhibiting overlapping or adjacent immunofluorescent staining as described in the text. The percentage of cilia-positive structures associated with centrosomes (γ-tubulin) or IFT88-positive cilia was calculated for each KV by dividing the number of associated cilia by the total number of cilia counted. Similarly, the percentage of γ-tubulin-positive centrosomes associated with Rootletin was obtained by dividing the number of colocalized centrosomes by the total number of centrosomes counted.

### Average cilia length

Cilia length was measured using maximum intensity projections of KV images captured at 12 hpf. Z-slice selection and Imaris segmentation were performed to isolate the KV region in each image. Within the segmented projection, a line was drawn along the central axis of each visible cilium using the FIJI/ImageJ line tool from the base to the tip. Measurements were recorded for all cilia in the KV, and the average cilia length per KV was calculated by dividing the total cilia length by the total number of cilia. Data were presented by averaging cilia lengths relative to the lumen area of the embryos from which they were quantified.

### Lumen area

Lumen area was quantified from *Tg(Sox17:GFP-CAAX)* and *Tg(Sox17:GFP)*, marking the KV region. Using FIJI/ImageJ tools, an ROI was drawn around the lumen boundary from a volumetric projected image of a 3D confocal data set of the KV. See previously published studies (Aljiboury et al., 2023; Rathbun et al., 2020; Wu et al., 2025).

### vEM sample preparation

Zebrafish embryos at 12 hpf were fixed in freshly prepared 4% PFA. The KV region was dissected and deyolked in 4% PFA, then oriented on a gridded Mattek dish using a fluorescence stereoscope. Post-fixation was performed in Karnovsky’s fixative (2.5% glutaraldehyde, 2% formaldehyde in 0.1 M sodium cacodylate, pH 7.4) for 2 hours at room temperature, followed by five 3-minute washes in 0.1 M sodium cacodylate buffer (pH 7.4). Samples were stored at 4°C until further processing. Samples were rendered stationary for subsequent steps by dropping a small but sufficient volume of 1% low-melt agarose solution directly on the gridded Mattek plate to embed the sample *in situ*. Samples were then incubated in 2% aqueous osmium tetroxide and 1.5% potassium ferricyanide for 1 hour, then rinsed in ddH₂O until the agarose was visibly clear. Samples were stained overnight at 4°C with 1% aqueous uranyl acetate and rinsed five times in ddH₂O. Lead aspartate staining was performed by incubating samples in a solution of 0.066 g lead nitrate in 10 mL of 0.03 M aspartic acid (pH adjusted to 5.5 with 1 M KOH) at 60°C for 30 minutes, followed by thorough rinsing in ddH₂O. Dehydration was carried out using a graded ethanol series (35%, 50%, 70%, 95%, and 100%),10 minutes at each step. Dehydration helped solidify the agarose-embedded sample to the point that it could be carefully moved into a glass vial. After this, samples were washed thrice for 10 minutes each in propylene oxide. Tissue infiltration was achieved with Polybed 812 resin using a graded sequence: 1:3 resin: propylene oxide for 1 hour, 1:1 resin: propylene oxide overnight, 3:1 resin: propylene oxide for 5 hours, and finally, 100% resin overnight. The agarose-embedded embryos were transferred to embedding molds containing freshly degassed resin and polymerized at 60–65°C for 48 hours. Embedded tissue blocks were trimmed into a pyramid shape, and the position of the KV was verified using X ray microscopy (Zeiss/Xradia Versa 730). The FIJI plugin Crosshair was used to target the precise angle and depth of the KV in the resin block based on the X ray microscopy data (Meechan et al., 2022). Ultrathin serial sections (∼70 nm) were cut with a diamond knife (Diatome) onto an ITO coverslip (SPI supplies) using a Leica ARTOS ultramicrotome. Coverslips were dried flat on a slide warmer set to ∼55°C for 30 minutes to ensure section adherence and drying.

### vEM image acquisition and reconstruction

Coverslips were mounted onto a 4-inch type-p silicon wafer (EMS) and grounded with conductive copper tape (EMS). Wafers were affixed to a 4-inch stage-decel holder in Ziess SEM (GeminiSEM 450; Carl Ziess) and imaged using ATLAS 5 vEM software (Fibics). Two ITO coverslips were imaged using a four-quadrant backscatter detector, with electron beam operated at 3.2 kV EHT with 2 kV beam deceleration and 600pA probe current. Low-resolution overview scans were collected at 3000 nm pixel resolution, medium-resolution section sets were collected at 150 nm pixel resolution, and high-resolution sites were collected at 10 nm pixel resolution. Once image acquisition was complete, the image stack was locally cropped and aligned using the ATLAS 5 software. The resulting image stack was exported and then processed using python-based scripts to produce an aligned and contrast/brightness adjusted image dataset: a stack of images at 10 nm (xy) x 100 nm (z). Higher resolution imaging of specific cilia was performed on the same ITO coverslips. The re-imaged areas were collected on the Ziess SEM using a four-quadrant backscatter detector, with electron beam operated at 3.5 kV EHT with 2 kV beam deceleration and 600 pA probes current. High resolution sites of targeted cilia were collected at 3 nm xy pixel resolution and exported. Original imaging notes were used to identify and re-image regions of interest.

### Manual segmentation in dragonfly software

3D segmentation of KV structures was performed manually using Dragonfly software. Image stacks, acquired from vEM, were imported into Dragonfly to create a volumetric dataset. Each structure of interest was then manually traced across serial sections using the software’s segmentation tools. The final reconstructed volume was generated by aligning individual segmented axis providing a detailed 3D representation of the sample.

### Statistical Analysis

PRISM9 software was used for all graph preparations that include all individual data points across embryos and clutches denoted by color and size of points respectively and as noted in legends. These plots were presented as bar graph with superplot to denote individual embryos across clutches (Lord et al., 2020). Unpaired two-tailed t-tests were performed using PRISM9 software. **** denotes a p-value<0.0001, *** p-value<0.001, **p-value<0.01, *p-value<0.05, n.s. not significant.

