## Supplementary material for "Functionally Essential and Structurally Diverse: Insights into the zebrafish Left-Right Organizer’s Cilia via Optogenetic IFT88 Perturbation and Volume Electron Microscopy": Table S1

**Table S1. Statistical analysis.**

| **Figure** | **Category** | **n embryo** | **n**  **Clutch** | | **Statistical Test** | **Parameters** | **Result** | | **p-value** |
| --- | --- | --- | --- | --- | --- | --- | --- | --- | --- |
| 1D | CRY2 | n=13 | n=3 | | Unpaired t test | t=3.113, df=23 | ** | | 0.0049 |
|  | Global IFT88 clustering | n=12 |  |  |  |  | ** | | 0.0049 |
|  | CRY2 Tg | n=22 |  |  |  | t=4.311, df=39 | *** | | 0.0001 |
|  | KV specific IFT88 clustering | n=24 |  |  |  |  | *** | | 0.0001 |
| 1E | CRY2 | n=13 | n=3 | | Unpaired t test | t=10.27, df=634 | **** | | <0.0001 |
|  | Global IFT88 clustering | n=12 |  |  |  |  | **** | | <0.0001 |
|  | CRY2 Tg | n=22 |  |  |  | t=19.81, df=1155 | **** | | <0.0001 |
|  | KV specific IFT88 clustering | n=24 |  |  |  |  | **** | | <0.0001 |
| 1F | CRY2 | n=13 | n=3 | | Unpaired t test | t=4.914, df=23 | **** | | <0.0001 |
|  | Global IFT88 clustering | n=12 |  |  |  |  | **** | | <0.0001 |
|  | CRY2 Tg | n=22 |  |  |  | t=6.433, df=39 | **** | | <0.0001 |
|  | KV specific IFT88 clustering | n=24 |  |  |  |  | **** | | <0.0001 |
| 2E | Ac-tub: Gamma-tub | n=29 | n=3 | | N/A | N/A | N/A | | N/A |
|  | Ac-tub: IFT88 | n=18 |  |  |  |  |  |  |  |
|  | Gamma-tub: Rootletin | n=21 |  |  |  |  |  |  |  |
| **Array tomography** | | | | | | | | | |
| **Figure** | **Category** | | | **n structure** | | | | **n embryo** | |
| 4E | Mother centriole plus daughter centriole | | | n=67 cilia | | | | n=1 | |
|  | Mother centriole | | |  |  |  |  |  |  |
|  | No centriole | | |  |  |  |  |  |  |
| 5C | Rootlet | | | n=59 centrosome | | | | n=1 | |
|  | DA | | |  |  |  |  |  |  |
|  | SDA | | |  |  |  |  |  |  |
|  | DA and SDA | | |  |  |  |  |  |  |
|  | CPMs | | |  |  |  |  |  |  |
|  | CaDR | | |  |  |  |  |  |  |
| 6H | CaDVs | | | n=67 cilia | | | | n=1 | |
|  | CaVs | | |  |  |  |  |  |  |
|  | Ciliary pocket | | |  |  |  |  |  |  |
|  | NIVB | | |  |  |  |  |  |  |
