## Supplementary material for "Functionally Essential and Structurally Diverse: Insights into the zebrafish Left-Right Organizer’s Cilia via Optogenetic IFT88 Perturbation and Volume Electron Microscopy": Table S2

**Table S2. SUPPLEMENTARY KEY RESOURCE TABLE**

| **Reagent or resource** | **Source** | **Identifier** |
| --- | --- | --- |
| **Antibodies** | | |
| Anti-IFT88 | Proteintech | 13967-1-AP; RRID: AB_2121979 |
| Rootletin | Fisher scientific | ABN1686MI |
| Acetylated Tubulin | Sigma Aldrich | T6793: RRID: AB_477585 |
| γ-tubulin | Sigma Aldrich | T5192; RRID: AB_261690 |
| Anti-GFP (Chicken) | GeneTex | GTX13970: AB_371416 |
| Anti-GFP (Rabbit) | Molecular Probes | A-11122: AB_221569 |
| Alexa Fluor Anti-Chicken 488 | Fisher scientific | A11039 |
| Alexa Fluor Anti-Rabbit 488 | Life Technologies | A21206; RRID: AB_2535792 |
| Alexa Fluor Anti-Rabbit 568 | Life Technologies | A10042; RRID: AB_2534017 |
| Alexa Fluor Anti-Rabbit 647 | Life Technologies | A31573; RRID: AB_2536183 |
| Alexa Fluor Anti-Mouse 488 | Life Technologies | A21202; RRID: AB_141607 |
| Alexa Fluor Anti-Mouse 568 | Life Technologies | A10037; RRID: AB_2534013 |
| Alexa Fluor Anti-Mouse 647 | Life Technologies | A31571; RRID: AB_162542 |
| **Chemicals, Peptides, and Recombinant Proteins** | | |
| DAPI | Sigma Aldrich | D9542-10mg |
| Agarose | Thermo Fischer | 16520100 |
| BSA | Fisher Scientific | BP1600-100 |
| BIO BASIC Maxi Prep Kit | BIO BASIC | 9K-0060023 |
| Dimethylsulphoxide | Fisher Scientific | BP231-100 |
| Paraformaldehyde | Fisher Scientific | O4042-500 |
| Phosphate Buffered Saline | Fisher Scientific | 10010023 |
| Molecular Probes Prolong Gold Antifade mount | Fisher Scientific | P36934 |
| Triton X-100 | Fisher Scientific | BP151500 |
| Tween 20 | ThermoFischer | BP337500 |
| Sodium Chloride | Fisher Scientific | BP358 |
| NEBuilder HiFi DNA assembly Cloning Kit | New England BioLabs | E5520S |
| mMESSAGE mMACHINETMSP6 | Invitrogen | AM1340 |
| OneTaq One-Step RT-PCR Kit | New England Biolabs | E5315S |
| **Reagent or resource** | **Source** | **Identifier** |
| Karnovsky’s fixative  20% Formaldehyde  10% Glutaraldehyde | EMS  EMS | 15713  16120 |
| Low-melt agarose | Millipore Sigma | A9045-10G |
| 4% Osmium tetroxide aqueous solution | EMS | 19170 |
| Sodium cacodylate trihydrate | EMS | 12310 |
| Potassium ferricyanide | EMS | 20150 |
| Uranyl acetate | EMS | 22400 |
| Lead aspartate solution  Lead nitrate  Aspartic acid | EMS  Millipore Sigma | 17900  A8949-25G |
| Ethanol | EMS | 15055 |
| Propylene oxide | Millipore Sigma | 110205-500ML |
| Polybed 812 resin | Polysciences  Polysciences  Polysciences  EMS | Poly/bed® 812 embedding media: 08791  Nadic Methyl Anhydride (NMA): 00886  Dodecenylsuccinic anhydride (DDSA): 00563  Benzyldimethylamine (BDMA): 11400 |
| **Equipment and consumables** | | |
| 35 mm Dish\| No.1.5. coverslip\| 20 mm Glass Diameter | MatTek Corporation | P35G-1.5-20-C |
| 40 mm x 22 mm No. 1 8-12 Ohm/sq ITO coverslip | SPI Supplies | 06497-AB |
| Silicon wafer | EMS | 71893-07 |
| Conductive copper tape | EMS | 77802 |
| 45 Ultra Diamond knife | Diatome | 25-US |
| ARTOS Ultramicrotome | Leica | https://www.leica-microsystems.com/products/sample-preparation-for-electron-microscopy/p/artos-3d/ |
| **Experimental models, organisms, and strains** | | |
| Zebrafish | Heidi Hehnly lab, Syracuse University | Tg(Sox17:cry2-GFP) |
| Zebrafish | Dasgupta and Amack, 2016 [1] | Tg (Sox17:GFP-CAAX)sny101 |
| Zebrafish | Navis et al., 2013 [2] | TgBAC(cftr-GFP) |
| Zebrafish | Zebrafish International Resource Center | Tg(sox17:GFP) |
| Zebrafish | Zebrafish International Resource Center | AB-Wildtype |
| **mRNA** | | |
| CRY2 | Rathbun et al., 2020 [4] | Plasmid: pCS2-CRY2; Addgene Plasmid #140572 |
| CIB1-RFP-IFT88 | This paper | Plasmid: pCS2- CIB1-RFP-IFT88 |
| **Software and algorithms** | | |
| 3D Dragonfly software | ORS Dragonfly | https://dragonfly.comet.tech/ |
| ATLAS 5 Array Tomography software (Fibics/Zeiss) | Fibics Inc | http://www.fibics.com/ |
| Python-based scripts | CCR volume EM | https://crtp.ccr.cancer.gov/vem/ |
| ImageJ/FIJI | NIH and Laboratory for Optical and Computational Instrumentation | https://imagej.net/Fiji |
| IMARIS, Bitplane | Oxford Instruments | https://imaris.oxinst.com/ |
| PRISM9 | GraphPad | https://www.graphpad.com/scientific-software/prism/ |
| LAS-X Software | Leica Microsystems | https://www.leica-microsystems.com/products/microscope-software/p/leica-las-x-ls/ |
| VisiView | Visitron | https://www.visitron.de/products/visiviewr-software.html |
